# Shifted levels of sleep and activity under darkness as mechanisms underlying ectoparasite resistance

**DOI:** 10.1101/2023.10.30.564749

**Authors:** Joshua B. Benoit, Oluwaseun M. Ajayi, Ashley Webster, Karl Grieshop, David Lewis, Hailie Talbott, Joy Bose, Michal Polak

## Abstract

Parasites harm host fitness and are pervasive agents of natural selection capable of driving the evolution of host resistance traits. Indeed, host resistance in natural populations typically shows ample genetic variation, which may be maintained when parasite resistance imposes fitness costs on the host in the absence of parasites. Previously we demonstrated significant evolutionary responses to artificial selection for increasing behavioral immunity to *Gamasodes queenslandicus* mites in replicate lines of *Drosophila melanogaster*. Here, we report transcriptional shifts in metabolic processes between selected and control fly lines based on RNA-seq analyses. We also show decreased starvation resistance and increased use of nutrient reserves in flies from mite-resistant lines. Additionally, resistant lines exhibited increased behavioral activity, reduced sleep, and elevated oxygen consumption under conditions of darkness. Using an independent panel of *D. melanogaster* genetic lines exhibiting variable sleep durations, we found a positive correlation between mite resistance and reduced sleep, providing additional support for a link between resistance and sleep. Experimentally restraining the activity of artificially selected mite-resistant flies during exposure to parasites under dark conditions reduced their resistance advantage relative to control flies. The results suggest that ectoparasite resistance in this system involves increased dark-condition activity and metabolic gene expression at the expense of nutrient reserves and starvation resistance.

**Significance statement:** Parasites are potent agents of selection, yet resistance may often be constrained evolutionarily because of trade-offs involving other fitness-related traits. Using artificial selection, we show that resistance to ectoparasites directly increases metabolism and decreases starvation resistance, predominantly through altered sleep and activity patterns at night. These studies highlight that active-resting patterns of the host are a significant driving force in ectoparasite resistance, but may have a negative impact on fitness during periods of low food availability. Our strongly integrative work suggests that parasite pressure may influence the evolution of host sleep and activity patterns.

## Introduction

Parasites are ubiquitous, and parasite-mediated selection can represent a strong evolutionary pressure that influences many host phenotypic traits and ecological processes (1–4). Defensive adaptations against parasites are common in plant, animal, and microbial host populations, where genetic variation in resistance is an essential component of host capacity to respond evolutionarily to parasite pressure (5–9). *Drosophila* and the many parasites that target species within this genus represent excellent systems to examine mechanisms of resistance and the factors that promote and constrain their evolution (10–14). Specifically, associations between *Drosophila* and ectoparasitic mites (15–19) are exceptional models for studies of ectoparasitism, as many such systems that are naturally occurring are also experimentally tractable, and the behavioral traits flies deploy against mites parallel those seen in other animal hosts (20). The benefits ectoparasitic mites may derive from attaching to flies include increased nutrient intake and reproductive output, and improved dispersal to new habitats (21). The host is harmed through damage to the cuticle and resource extraction along with costly immune and repair responses (5, 15, 18, 22). The combined effects of mite-derived damage negatively influence the expression of male and female reproductive tissues, mating success, and lifespan (5, 15, 22). Other host behavioral and physiological changes that occur in response to parasitism or parasite exposure include adaptive shifts in male reproductive effort (23), anti-mite defensive behaviors (24), oviposition site preference (25), and rates of respiration (26).

*Drosophila melanogaster* and other fruit fly species have been noted to be parasitized by *Gamasodes* mites in Asia (Taiwan and Thailand), and Australia (17, 19, 22). The extended geographic distribution and host range of *Gamasodes* mites indicates that they are likely to be a pervasive selective force affecting many aspects of fly biology. The behaviors that flies use to avoid infestation by ectoparasitic mites include bursts of flight from the substrate, locomotor responses, and vigorous grooming once the mite has made contact with the fly (24, 27). Whereas strong selection is likely to act on these first-line forms of defense (27, 28), the molecular and physiological bases of these defensive traits are yet to be established.

In the present study of *D. melanogaster*, RNA-seq and functional studies revealed that metabolic activity increased in response to artificial selection for increased behavioral resistance to *Gamasodes* mites (27). The increased metabolism was associated with reductions in nutrient reserves and starvation resistance and was correlated with increased activity and reduced sleep during the dark. These results identify metabolic and activity changes as a potential causal factor underlying ectoparasite resistance. Resistance assays of an independent panel of *Drosophila* genetic lines with variable sleep supported a link between decreased sleep and enhanced parasite avoidance. The results suggest that increased activity and reduced sleep could be a critical factor in preventing ectoparasite attachment and that this energetically costly behavioral immunity trades off with nutrient reserves and starvation resistance. This study enhances our understanding of fundamental mechanisms that maintain genetic variation mediating a naturally occurring host-parasite symbiosis.

## Results

### Increased resistance coincided with transcriptional changes associated with metabolism

Previously we performed artificial selection for increasing resistance to *Gamasodes queenslandicus* mites in male *D. melanogaster* for 16 generations (27), which we continued in the present study for a total of 22 generations. The selection program resulted in significant evolutionary responses, yielding three replicate selection lines that were significantly more behaviorally resistant to ectoparasitic mites than their unselected counterparts (Fig. 1A). At the terminus of the 22 generations of selection, transcriptional analyses revealed that 108 genes were significantly differentially expressed (68 up-regulated and 40 down-regulated) in selection lines relative to control lines (three biological replicates of each of the three replicate lines per selection treatment) (Table S1). Gene ontology enrichment analyses revealed increased oxidoreductase activity in selection lines relative to controls along with increases in other factors, such as three genes involved in glycolysis (aldolase, acetyl-CoA synthetase, and phosphoglycerate kinase) (Fig. 1B, Table S1). These results suggest that there are potential differences in metabolism that occur in flies as a result of selection for mite resistance.

**Figure 1.**
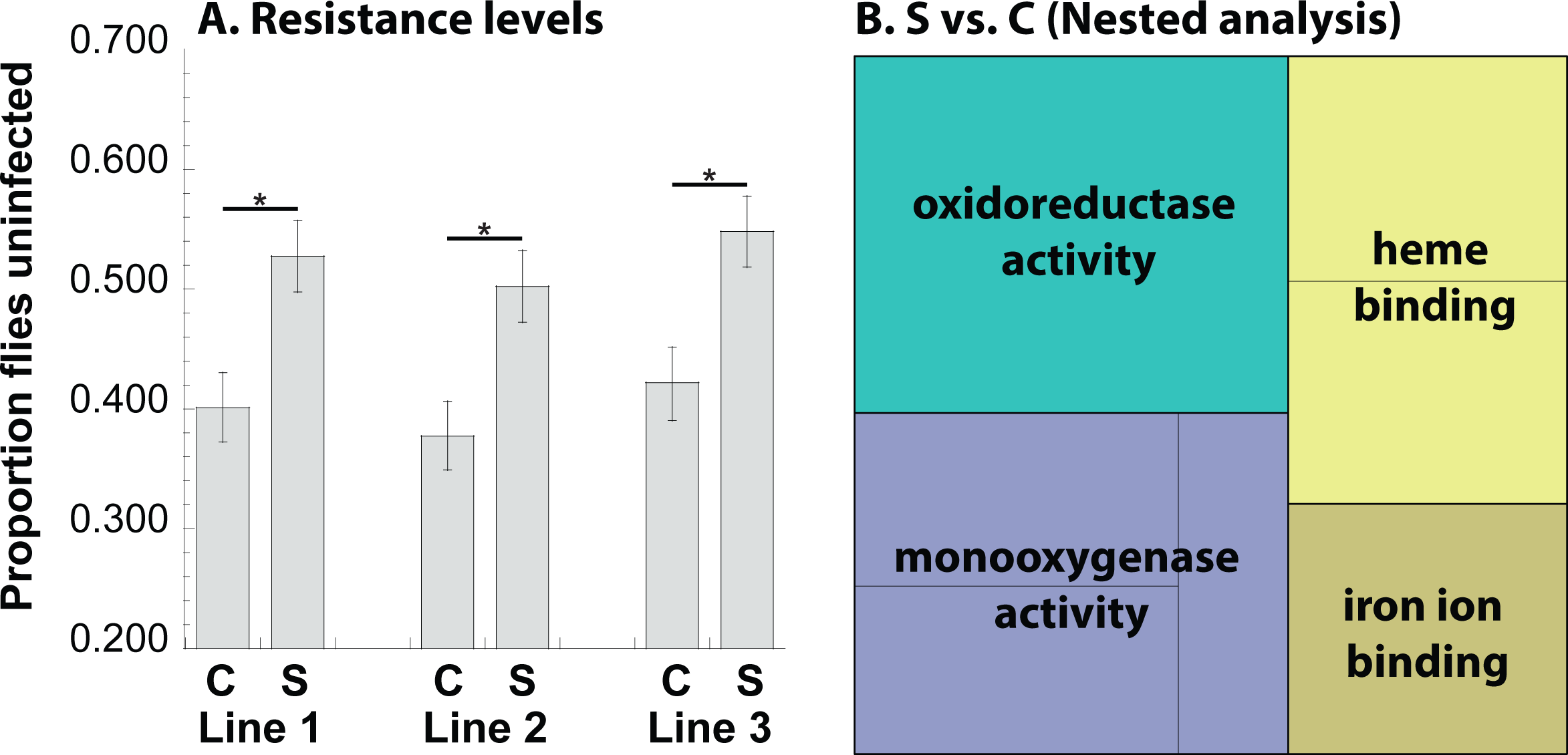
Selection increases resistance to mites and RNA-seq studies reveal altered metabolic-associated factors following selection for ectoparasite resistance. A. Artificial selection for mite resistance resulted in significantly reduced susceptibility to mites in selected (S) versus control (C) lines after 21 generations. * denotes significance at P < 0.05 between control and selected lines. (F = 16.24, d.f. = 2,5, P = 0.001) Error bars represent the 95% confidence intervals. B. Gene ontology categories enriched in differentially expressed genes between S and C from a full analysis of all samples (nested within the replicate, revealing factors generally associated with metabolism, FDR, P < 0.05). There were three biological replicates of RNA-Seq data for each treatment (S and C) of the three artificial selection replicates.

### Starvation resistance decreased with increased metabolism under dark periods

Compared to control lines, the observed upregulation of metabolism-associated factors among resistant lines suggests that a trade-off exists between resistance and host nutrient reserve management. Consistent with this expectation, starvation resistance in ectoparasite-resistant lines was reduced compared to control lines (Fig. 2A). This apparent trade-off was corroborated by the observation of greater reductions in body lipid and protein reserves after 36 hours of starvation in resistant lines relative to controls (Fig. 2B, C). This difference in levels of nutrient stores suggests relatively increased lipolysis and proteolysis in the resistant lines. Thus, there are discernible metabolic phenotypes that co-evolved with ectoparasite resistance under artificial selection, identifying potential cost of ectoparasite resistance as a reduction in host survival resulting from accelerated reduction in body condition under nutrient deprivation.

**Figure 2.**
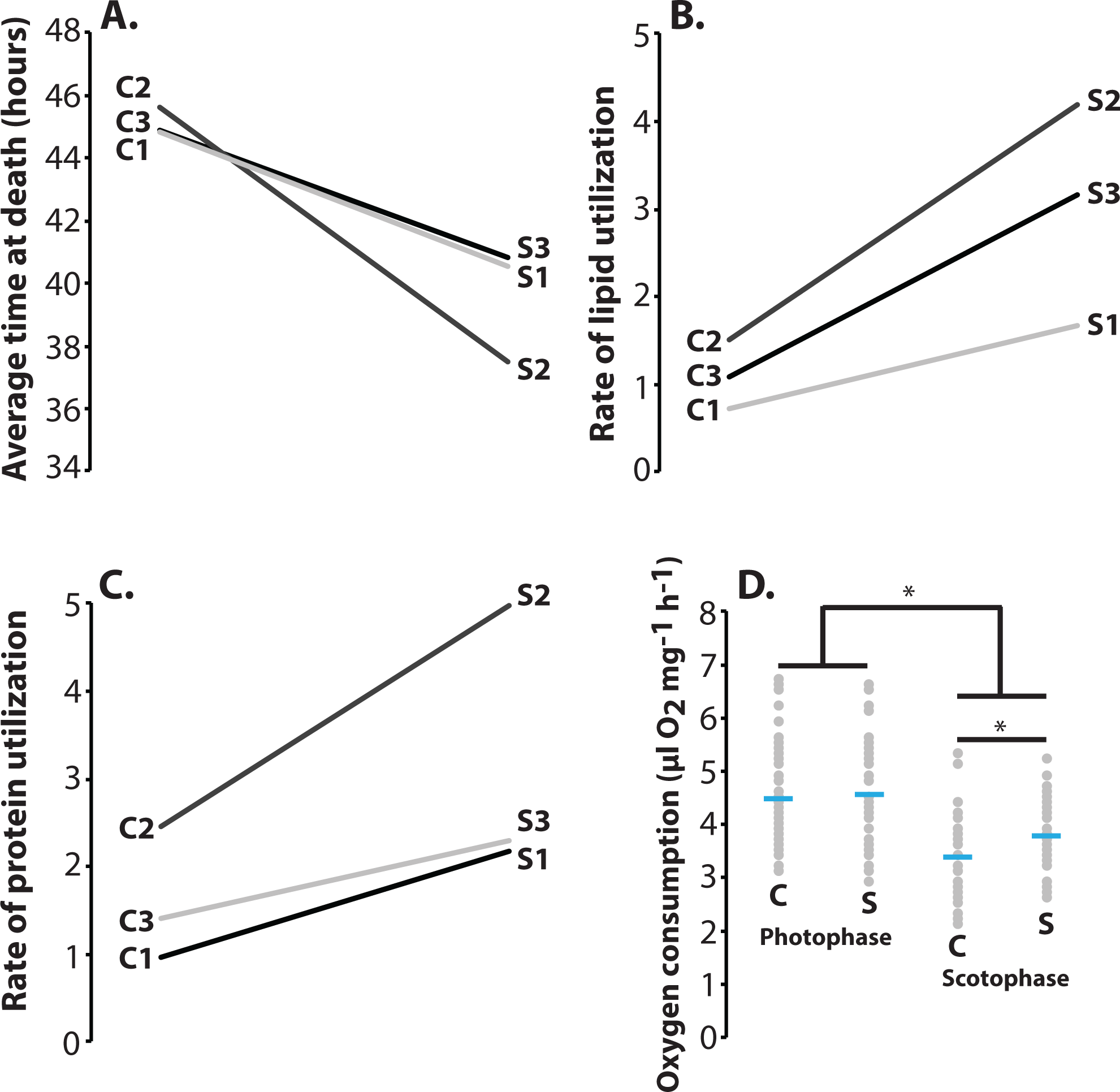
Flies with increased resistance have higher metabolism during the dark conditions. A. Death due to starvation was increased following selection in all three lines. Kaplan-Meier survival analysis, generalized Wilcoxon test, Chi-square = 18.234, P < 0.001 for control compared to selected for each line, N = 150 individuals per line. B and C. Lipolysis (F = 11.84, d.f. = 2,5, P = 0.031) and protein (F = 11.08, d.f. = 2,5, P = 0.001) breakdown was increased following selection in all lines. N = 3 groups of three flies examined per line. D. Oxygen consumption was increased in the selected flies under darkness (scotophase) compared to light conditions (photophase) (F = 10.47, d.f. = 1,4, P = 0.009). N = 15 flies measured per line when selected and control lines are compared without light. * denotes significance at P < 0.05 between control and selected lines.

To further probe this potential metabolic trade-off, we measured oxygen consumption of flies during a daily cycle of light and darkness. Surprisingly, oxygen consumption was not different between control and selected lines under light conditions (Fig. 2D). However, when oxygen consumption was measured in the dark, it was significantly higher in the resistant lines compared to controls (Fig. 2D). This result indicates that the metabolic costs of ectoparasite resistance may primarily manifest under dark conditions.

### Activity during darkness was greater in ectoparasite-resistant lines

As oxygen consumption was increased under dark conditions, we tested whether the fly activity and sleep profiles were altered following selection throughout a daily cycle of light and dark conditions (Fig. 3A). As with oxygen consumption, activity and sleep levels were not significantly different during the light. However, resistant lines did show a significant increase in activity and a decrease in the duration of sleep bouts during the dark relative to controls (Fig. 3B, C) – two traits that are certainly related but not necessarily one in the same. To establish if reduced sleep and increased activity profiles were associated with resistance to ectoparasitism, we exposed an independent panel of inbred fly lines that differed in activity and sleep levels to ectoparasitic mites (29). There was a significant relationship between the level of sleep under dark conditions and the proportion of flies parasitized by mites (Fig. 3D). This result supports the possibility that sleep and activity levels are causal factors affecting ectoparasite resistance in flies.

**Figure 3.**
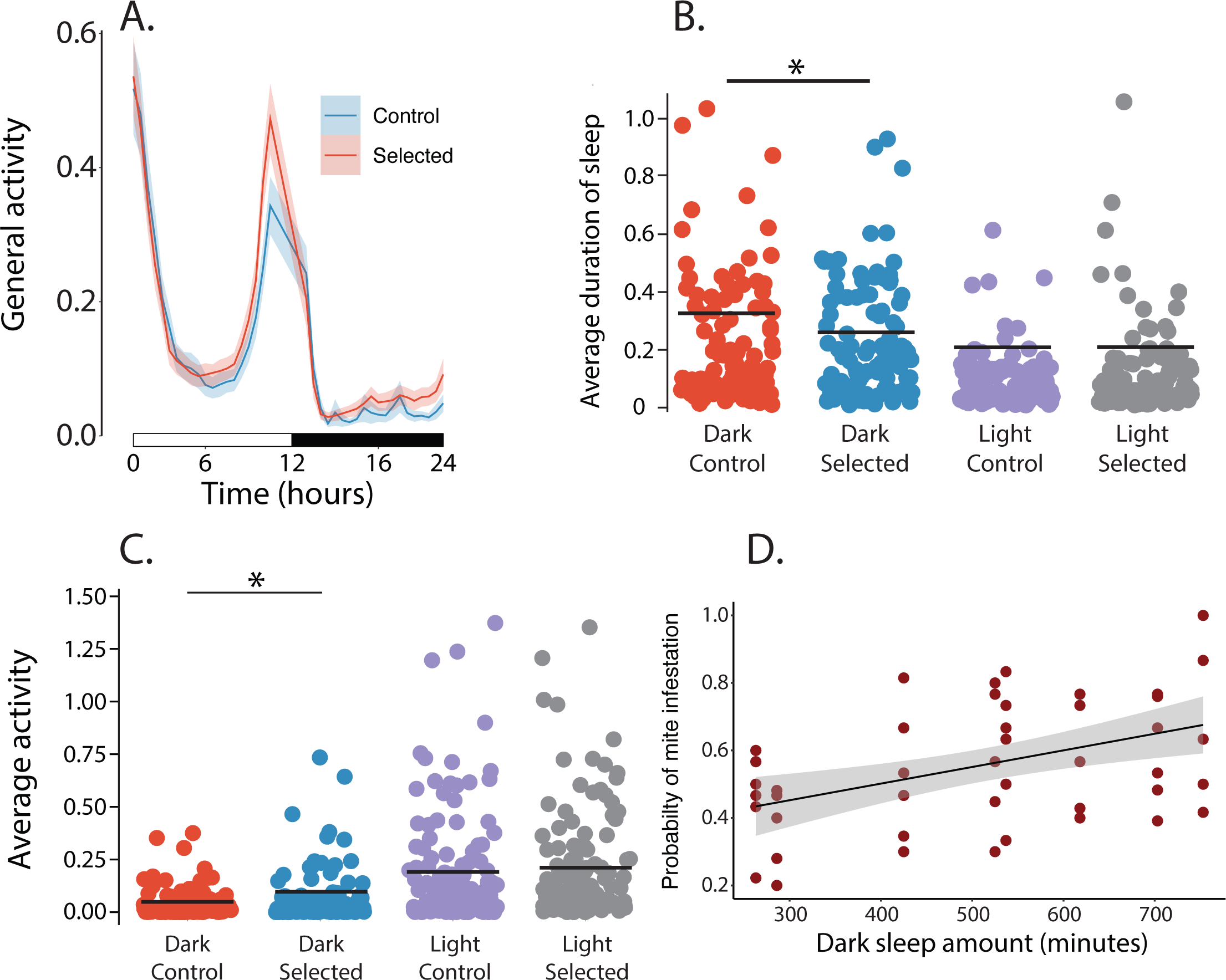
Sleep and activity profiles are altered in selected lines. A. Daily activity profile for control and selected fly lines, all lines combined. Shaded areas represent the 95% confidence intervals. N = 40-45 individuals per line. B. Sleep duration decreased during the dark cycle for lines selected for resistance (F = 19.291, d.f. = 1,4, P < 0.001). Other aspects of sleep, such as bouts or duration of bouts, are not significantly different between selected and control lines ( P > 0.05 in all cases). C. Activity is increased during the night for lines selected for resistance (F = 15.98, d.f. = 1,4, P = 0.002).Each point represents the activity of a single fly. D. A significant interaction occurs between an independent panel of eight inbred lines with different sleep levels (29) and susceptibility to mites (F = 3.021, d.f. = 7,40, P = 0.012). N = 6 for each line was assayed for parasitism. * denotes significance at P < 0.05 between control and selected lines.

### Movement restriction and assay under light conditions reduced ectoparasite resistance

To confirm in another way the causal role of activity and sleep-like behavior in mediating ectoparasite resistance, we manipulated activity by restraining flies. We found that selected lines no longer showed improved under darkness resistance relative to controls when restrained (Fig. 4A), matching a previous study showing that flight restriction erodes mite resistance (27). Secondly, we performed mite parasitism assays under light and dark conditions, and showed that resistance divergence between selected and control lines was reduced in light relative to dark conditions (Fig. 4B). These results support the conclusion that increased activity under dark conditions and reduced sleep-like behavior are important factors mediating host response to selection for improved mite resistance.

**Figure 4.**
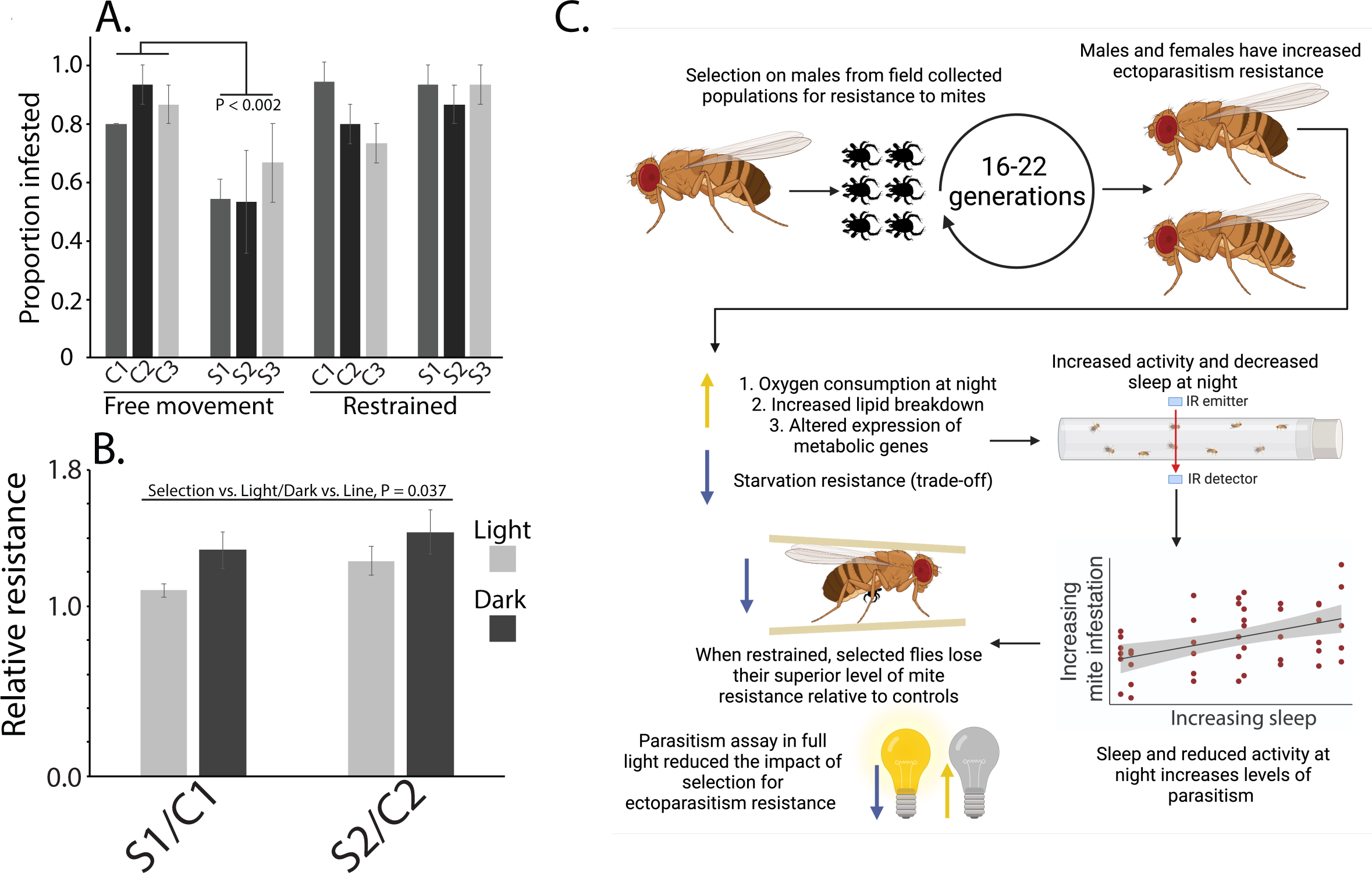
Factors that alter sleep status shift ectoparasite resistance. A. Restraining flies reduces the resistance of selected lines compared to controls, (F = 28.66, d.f. = 1,4, P < 0.002). B. The effect of selection on mite resistance is more profound when flies are exposed to mites under dark conditions. N = 4 parasitism assays per line (F = 11.839, d.f. = 1,3, P = 0.016). C. Summary of dynamics between sleep and activity in relation to resistance to ectoparasitic mites. This highlights that altered sleep and activity patterns are associated with resistance to parasitism.

## Discussion

This study indicates that declining sleep and increased activity are critical mechanisms for preventing ectoparasitism in fruit flies. Following artificial selection for resistance (27), specific metabolic transcriptional pathways associated with increased monooxygenase activity along with a multitude of other factors such as specific genes underlying glycolysis (*e.g.* aldolase, acetyl-CoA synthetase, and phosphoglycerate kinase) were up-regulated. Metabolic activity increased at the expense of starvation resistance in flies resistant to mite parasitism. This finding matched increased activity and reduced sleep during dark conditions in selected versus control flies, which is one likely factor governing the evolution resistance to ectoparasitic mites. We provided experimental confirmation that altered sleep and activity profiles are critical for ectoparasite resistance in experiments where fly activity was prevented and by using fly lines expressing variable sleep levels.

Sleep and host-parasite dynamics have been studied in multiple systems, but these have predominantly involved endoparasites such as malaria (30, 31). In general, the response to internal parasites requires a robust immune response, where increased sleep may improve this response by allowing the increased expression of immune factors (30, 31). In contrast, host sleep (or lack thereof) could influence ectoparasite resistance differently, as behavioral resistance, rather than immune response, is of greater importance to avoid infestation. Here, we confirmed that among resistant lines, sleep decreased and activity increased during darkness, which is when sleep predominantly occurs in *D. melanogaste*r (32, 33). The pattern of reduced sleep likely allows the flies to be more responsive during encounters with mites under dark conditions. There could be differences in sleep occurring during light conditions, but these differences may be less important as *D. melanogaste*r sleeps much less during the day (32, 33). Interestingly, as the artificial selection was conducted under both light and dark conditions (spanning the diurnal transition) (27), this raises the possibility that a small change in sleep patterns or activity in the dark could have been a direct target of selection that yielded the increased resistance to mite infestation. The altered sleep and activity profiles are likely to be a general phenomenon that occurs among animals in the presence of ectoparasites. Parasitism has been noted in birds and bats to be associated with shifts in sleep and rest behavior during nest cleaning and grooming (34, 35). Changes in sleeping patterns in primates have also been suggested to reduce exposure to ectoparasites (36). The present study is unique in revealing a potential causal link between reduced host sleep, increased activity, and improved resistance to ectoparasites. Recently, phototaxis effects have been noted during fly-mite interactions (37), where *Drosophila nigrospiracula* exposed to parasitic mites under light conditions had lower mite parasitism than those held in the dark with mites. Furthermore, flies will reside more in lighted areas after exposure to mites (37). As light exposure increases fruit fly activity and reduces sleep (32, 33), the studies on *D. nigrospiracula* provide additional support for our observations in *D. melanogaster* that activity and sleep influence fly-mite interactions.

Parasite resistance may be associated with distinct fitness costs, which include reduced stress tolerance and damaged fecundity identified in multiple animal systems (2–4) including other fly-mite systems (5, 27). The increased expression of metabolism-associated genes and altered starvation resistance in selection lines reported here suggest that mite resistance is likely to alter fly physiology in relation to metabolic output, which we demonstrated through the assessment of lipid reserves during starvation, and altered oxygen consumption. Our artificial selection experiment may have allowed mite resistance to evolve despite these metabolic costs because the *ad libitum* food conditions reduced the strength of selection for nutrient management. Importantly, direct parasitism by mites induces a physiologically costly immune response (5, 22), and significantly reduces host egg production and lifespan (23, 38), thus avoiding parasitism is critical. Thus, the optimal degree of metabolically demanding mite resistance behaviors in nature is likely determined by the balance of parasite pressure versus resource limitation and nutrient management, which could vary temporally, geographically and taxonomically.

The finding that altered sleep patterns impact ectoparasite resistance represents a novel form of host resistance, similar to those involved in predator-prey dynamics. Indeed, many ectoparasites such as blood-feeding flies and mites have been classified as micropredators (39, 40), and differences in sleep have been associated with predation levels (41–43) such that animals with increased exposure to predators have reduced or lighter sleep (42, 44). Here, we showed that reduced sleep-like behavior in *D. melanogaster* was the target of selection for parasite resistance, providing evidence that sleep behavior can (co-)evolve as a result of interactions with an ectoparasite. The phenotypes confer ectoparasite resistance with a direct metabolic cost that could act as a trade-off constraining the evolution of resistance to the mites, especially when parasitism rates are low and/or nutrients limited.

## Acknowledgments

The research was supported by National Science Foundation (NSF) grant 1654417 (to M.P. and J.B.B.). Partial funding for reusable equipment was provided indirectly by the United States Department of Agriculture 2018-67013 and the National Institute of Allergy and Infectious Diseases, award numbers R01AI148551 and R21AI166633 to J.B.B. We thank Rachel Black and Hannah Tran for their excellent assistance in media preparation.

## Materials and methods

### Flies, mites, and selection for resistance

The flies (*Drosophila melanogaster* Meigen) and mites (*Gamasodes queenslandicus* Halliday and Walter) were collected at Cape Tribulation, Queensland, Australia (27). The artificial selection protocol for increasing resistance to mites, which is described in detail elsewhere (24, 27) in significantly increased resistance in response to 16 generations of sustained selection. At generation 16, strongly significant and consistent divergence between selected lines and their counterpart unselected control lines was observed (27, 45)not exceeding 22 generations of selection. Even though selection was only conducted on males, both sexes showed increase resistance (27). Furthermore, increased cross-resistance to another mite species (*Macrocheles subbadius* Berlese) was shown (27), indicating that the defensive mechanisms that were selected are likely general to ectoparasitic mites. Lines of *D. melanogaster* with varying levels of sleep were acquired from the Bloomington Stock Center from short and long sleep lines that were cross-bred to increase the levels of sleep (29). Male flies were used in these analyses.

### Mite infestation experiments

Mite infestation experiments were conducted according to previously developed methods (5, 22, 24, 38) that allow for determining infestation differences between fly lines. These were assays similar to those used for selection, where flies were exposed to mites in a jar containing media used to rear *G*. *queenslandicus* (22, 24, 38). Each assay measured ectoparasitism in each fly line relative to its paired control line across a set of replicate infestation chambers. For each comparison, flies equal in size were aspirated into a chamber (total flies ranged between 40 and 50 flies). Minute wing clips at the tip of each wing were used to distinguish groups (≤ 3% of the wing), which have been previously shown to have no impact on fly susceptibility to mites (22, 24), and clipping was alternated between groups across chambers. A recovery period of 24h was used after clipping the wing under CO_2_. These jars were examined after 6-8 hours based on the relative number of mites in the media. The Direct presence of mites and scars on flies from chambers in relation to the wing clipping that marks the specific lines was used to assess the prevalence of parasitism. These parasitic assays were conducted on the selected and control lines after 21 generations under either full light or in complete darkness (selection line S3 was lost during COVID-19 restrictions as such only S1 and S2 were used in the light and dark comparative analyses) and for flies from the sleep inbred panel. For the sleep inbred panel, lines from the panel were intermixed during the assays to allow for random comparisons between each line.

For susceptibility tests under restraint, flies were transferred to shortened 200-µl micropipette tips, a confined space that severely limits fly movement (45, 46), and exposed to two mites, effectively mimicking periods of sharply reduced activity. After two hours in the dark, the flies were removed and examined for attached mites or mite-induced scars. This assay was conducted on the selected and control lines after 20 generations of artificial selection.

### RNA-seq analysis following selection

Flies were collected after 16 generations of selection for the RNA-seq studies. RNA was extracted from the flies as previously described (22) in three biological replicates per line. Illumina sequencing was conducted at Novogene on an Illumina NovaSeq Hiseq platform. Illumina datasets are available at the National Center for Biotechnology Information’s Sequence Read Archive: Bioproject PRJNA999910. Differential expression of genes was determined based on methods previously described (22, 47, 48). Briefly, the quality and presence of adapters were determined with FASTQC (49), followed by a filtering/trimming procedure with Trimmomatic v.0.36 using default settings (50). At least 30 million reads were available for each sample following trimming. Reads were mapped to the transcript assembly release 6.10 from Flybase (51) using Kallisto v0.48.0 with recommended settings (52) summarized to the gene level using tximport v1.12.3 (53). Differential gene expression analysis was performed with DESeq2 (54) with the read counts generated from Kallisto. Filtering in DESeq2 was conducted to remove predicted transcripts with fewer two mapped reads. Selection-specific effects to identify overlapping differences among lines were determined by a nested analysis that included each line and replicates. An adjusted p-value of ≤ 0.01 was used to identify genes as differentially expressed. Specific enriched gene ontology (GO) categories were determined using g:Profiler with gene sets increased and decreased during selection analyzed separately (55) and treemaps to visualize the GO was generated with REVIGO (56).

### Starvation assay, nutritional reserve levels, oxygen consumption

Flies were collected following 18-22 generations of selection for these different assays. Flies were transferred to 1% agar vials to provide water without nutrients (1 or 3 males per vial, no difference was noted in the duration of starvation between the two groups, and the results were combined), and the number of dead flies was noted every eight hours until all flies were dead. Nutrient reserve levels were measured during starvation based on previously used methods (22, 57), which use spectrophotometric assays to establish the lipid and protein levels in relation to the dry mass of flies and relative to a protein or lipid standard. Differences in oxygen consumption between select and control lines were determined using a microrespirometer (57, 58). In brief, individual flies were placed in microrespirometers were positioned in a water bath (25°C) and were allowed 15 minutes to equilibrate before measurements. CO_2_ production, measured through the movement of KOH, was measured every hour for 6 hours. Assuming that one mole of O_2_ is consumed for every mole of CO_2_ released, oxygen consumption was calculated using the distance traveled by the KOH and expressed as μl O2 mg dry mass^-1^ h^-1^.

### Sleep and activity assessment

The rest-activity rhythms of the flies (control, selected, and sleep lines) were quantified with the aid of a Locomotor Activity Monitor 25 (LAM25) system (TriKinetics Inc., Waltham, MA, USA) and the DAMSystem3 Data Collection Software (TriKinetics). Flies were collected after 19-22 generations of selection and those from the sleep inbred panel (29) were used for these assays. Individuals were loaded into 25 x 150 mm clear glass tubes with access to fly media provided *ad libitum*. The glass tubes were then positioned horizontally in the LAM25 system, allowing the simultaneous recording of 32 individuals in an “8 x 4” horizontal by vertical matrix. Replicates were intermixed between trials to allow for randomization. The entire set-up was placed in a light-proof incubator supplied with its own lighting system at 24°C, 70-75% RH, under a 12hr:12hr L/D cycle. Following the acclimation of individuals for 48 hours, activity level was measured in 1 min bins (as the number of times an individual crosses an infrared beam), and sleep levels were assessed as 5 min of continuous inactivity (29, 32). This allows for the determination of sleep bouts, duration of bouts, and total sleep. No differences were noted in the sleep bouts or total sleep, but the duration of bouts were significantly different between control and selected lines. Although sleep is related to activity, there are distinct differences in the quantification of activity and sleep, and sleep levels are not directly an inverse of general activity levels. Data collected with the DAMSystem3 was processed using the Rethomics platform in R with its associated packages including *behavr*, *ggetho*, *damr* and *sleepr* (59).

### Statistical analyses

For all analyses, we used the R software (60) and plots were generated with *ggplot*. Survival analyses were determined with a Kaplan-Meier test and compared with a general Wilcoxon test. General linear models were used to examine the interactions between selection status and response variables where replicates for each selection and control lines are nested as a random effect within the selection and control treatments to establish that effects are significant among control and selected groups. Statistical outputs and replicate numbers are provided within the figure legends.

